# Influence of myosin regulatory light chain and myosin light chain kinase on hair cells of the inner ear

**DOI:** 10.1101/2023.10.14.562357

**Authors:** Ryohei Oya, Kwang Min Woo, Brian Fabella, R. G. Alonso, A. J. Hudspeth

## Abstract

In the receptor organs of the inner ear and lateral line, sensory hair cells detect mechanical stimuli such as sounds, accelerations, and water movements. In each instance a stimulus deflects the hair bundle, a hair cell’s mechanically sensitive organelle. The bundle pivots upon the cell’s apical surface, which includes an actin meshwork called the cuticular plate and is surrounded by a ring of filamentous actin and non-muscle myosin II (NM2). Myosin regulatory light chain (RLC) is expressed at the apical surfaces of hair cells and RLC is additionally found in hair bundles. NM2 and the phosphorylation of RLC by myosin light chain kinase (MLCK) have earlier been shown to regulate the sizes and shapes of hair cells’ apical surfaces. We have found that inhibitors of NM2 and MLCK reduce the stiffness of hair bundles from the bullfrog’s sacculus. Moreover, MLCK inhibition inhibits the spontaneous oscillation of hair bundles and increases the resting open probability of transduction channels. In addition, MLCK inhibition elevates hearing thresholds in mice. We conclude that NM2 and the phosphorylation of RLC modulate slow adaptation and thereby help to set the normal operating conditions of hair bundles.

**Statement of significance:** Sensory hair cells play a key role in mechanoelectrical transduction by the inner ear and lateral-line system. To detect stimuli such as sounds and accelerations, hair cells use an active process to amplify their mechanical inputs. Although amplification is accomplished in part by the activity of the mechanosensitive hair bundles, the molecular mechanism of the active process remains uncertain. The present study shows that non-muscle myosin II (NM2) and the phosphorylation of myosin regulatory right chains (RLC) by myosin light chain kinase (MLCK) regulate the stiffness and spontaneous oscillation of hair bundles as well as the resting open probability of mechanotransduction channels. The results implicate myosin motors in the control of the active process of hair cells.

## Introduction

Sensory hair cells mediate mechanoelectrical transduction in the internal ear and lateral-line system. The site of mechanical sensitivity, which is conserved in the receptor organs of all chordates, is the hair bundle: an erect cluster of actin-filled rods, the stereocilia, protruding from the apical cellular surface. Graded in length across a hair cell’s surface, the stereocilia are conjoined by filamentous tip links that extend from each shorter stereocilium to the longest adjacent one (1). The lower end of each link is associated with two or more transduction channels that open in response to increased tension in the link and initiate an electrical response in the cell.

Myosin motors play several important roles in the transduction process of a hair cell. For example, a slow adaptation process, which typically exhibits a time constant of 10-30 ms, ensures that the cell’s response does not saturate in response to large stimuli (1). The insertional plaque at the upper end of each tip link, also known as the upper tip-link density, includes the proteins harmonin, vezatin, and SANS as well as myosin molecules (2,3): myosin 1c (MYO1C) in the frog’s saccule and mammalian vestibular organs (4), and myosin 7a (MYO7A) as well as MYO1C in the mammalian cochlea (5,6). In the absence of stimulation, the climbing activity of the myosin molecules maintains in each tip link a tension that biases the channels to a certain resting open probability (6–8). Moreover, after the bundle has been displaced in the excitatory or inhibitory direction, the myosin molecules at the tip link’s insertion respectively descend or climb to restore an approximately constant tension (9).

Although the elegance, regularity, and complexity of the hair bundle have attracted extensive attention from sensory biophysicists and developmental biologists, the cytoskeletal foundation on which the bundle rests is also of interest. The base of each stereocilium extends an actin rootlet that is anchored in the cuticular plate, a feltwork of actin filaments and spectrin (10). The cuticular plate is in turn stabilized by various structures. On the apical surface of a cochlear outer hair cell, the cuticular plate is surrounded by a circumferential ring of filamentous actin and non-muscle myosin II (NM2) (11,12). The interaction between these proteins mediates contraction of the apical cellular surface. Apical constriction of a similar form occurs in several developmental contexts, for example in the neural epithelium during closure of the neural tube (13,14) and in hair-cell morphogenesis (15), including the developmental alignment of hair bundles (16).

Like other myosin II molecules, NM2 consists of three pairs of proteins: two molecules of a myosin heavy chain (MYH), two of an essential light chain (ELC), and two of a regulatory light chain (RLC). The activity of NM2 is regulated through phosphorylation of the RLCs by myosin light chain kinase (MLCK) (12). When activated by the binding of the Ca^2+^-calmodulin complex, MLCK phosphorylates RLC (17), which in turn causes myosin heads to swing toward actin filaments, increases the duty ratio of myosin-actin interaction, and promotes force generation (18). Both RLC and MLCK occur in distinct isoforms in various cell types; for example, the organ of Corti expresses RLCs such as MYL9 and MYL12 as well as MYLK1 (also known as smooth-muscle MLCK or smMLCK). The phosphorylation of MYL12 by MYLK1 regulates shape changes in outer hair cells (12); moreover, deletion of the gene *MYLK1* in cochlear hair cells causes hearing loss (19,20).

Not only NM2 but also isoforms such as MYO1C and MYO7A can potentially bind RLC (21,22). As a result, the phosphorylation of RLC by MLCK might regulate several aspects of cellular shape and hair-bundle motility. In the present study, we therefore sought to determine the physiological consequences of pharmacological inactivation of NM2 and MLCK.

## Materials and Methods

### Animals

All animal experiments were performed in accordance with institutional guidelines provided by the Institutional Animal Care and Use Committee (IACUC) at The Rockefeller University. Adult American bullfrogs (*Rana catesbeiana*) of both sexes were purchased from Frog Pharm (Twin Falls, ID, USA). Female C57BL/6N mice of seven to eight weeks of age were obtained from Charles River Laboratories (Wilmington, MA, USA).

### Antibodies and reagents

We used the following antisera and dilutions: anti-MYH9 (ab238131; Abcam, Cambridge, UK; 1:50); anti-MYH10 (ab230823; Abcam; 1:50); anti-MYH14 (Proteintech, Rosemont, IL, USA; 1:50); anti-MYL12A/B (sc-376606; Santa Cruz, Dallas, TX, USA; 1:50); anti-MYL9 (ab191393; Abcam; 1:50); anti-phospho-myosin light chain 2 (3671; Cell Signaling Technology, Danvers, MA, USA; 1:50); Alexa Fluor 488-labeled secondary antibody (Thermo Fisher Scientific, Waltham, MA, USA; 1:500). F-actin was labeled with rhodamine-phalloidin (PHDR1; Cytoskeleton, Inc., Denver, CO, USA; 1:167).

We also employed the following reagents: Dako Protein Block Serum-Free (Agilent, Santa Clara, CA, USA); Dako Antibody Diluent (Agilent); blebbistatin (ab120425; Abcam); protease XXIV (Sigma-Aldrich, St. Louis, MO, USA); PeriAcryl90 (GluStitch Inc., Delta, BC, Canada); ML7 (ab120848; Abcam); MLCK inhibitor peptide 18 (19181; Cayman Chemicals, Ann Arbor, MI, USA).

### Immunofluorescence labeling

Samples were fixed for 30 min at room temperature with 4% formaldehyde in 0.1□M phosphate-buffered saline solution (PBS). After permeabilization for 30 min at room temperature in 0.2 % Triton X-100 in PBS, they were blocked for 30 min at room temperature with Dako Protein Block Serum-Free. The samples were incubated overnight at 4 □ with primary antisera in Dako Antibody Diluent. Sections were incubated for 60 min at room temperature in Alexa Fluor secondary antiserum in blocking solution. To label F-actin, rhodamine phalloidin was incubated in blocking solution for 60 min at room temperature. Confocal images were acquired with an IX83 laser scanning microscope (Olympus, Tokyo, Japan) with a UPlanSApo 60X objective lens (Olympus).

### Physiological recording from bullfrog saccules

The frog saccule was dissected as previously described (23). The dissection was performed in oxygenated artificial perilymph containing 114 mM Na^+^, 2 mM K^+^, 2 mM Ca^2+^, 118 mM Cl^-^, 3 mM **D**-glucose, and 5 mM HEPES at pH 7.3. The preparation was exposed for 35 min at 26 □ to 66 μg/mL protease XXIV in artificial perilymph. After the otolithic membrane had been detached from each specimen by the enzyme treatment, it was removed with fine forceps.

The previously reported protocol to build a two-compartment experimental chamber was optimized for the present study (23,24). The saccule was sealed with PeriAcryl 90 glue in a hole with a 500-700 μm diameter of a 10-mm square aluminum foil with the hair bundles facing upward. Vacuum grease was put on the surface of the lower and upper chambers to prevent the solutions from leaking and mixing with each other. Then, the foil-mounted saccule was placed on the lower chamber which was filled with the artificial perilymph. The upper chamber was placed on the foil and filled with artificial endolymph containing 2 mM Na^+^, 118 mM K^+^, 0.25 mM Ca^2+^, 118 mM Cl^-^, 3 mM **D**-glucose, and 5 mM HEPES at pH 7.3. To adjust the temperature, a temperature controller (TC200; Thorlabs, Newton, NJ, USA) was placed in the upper chamber.

The optical setting which was previously reported (25) was optimized for the present study. The frog saccule in a two-compartment chamber was placed under an upright microscope (BX51WI; Olympus, Center Valley, PA, USA) and imaged to visualize with a 60X water-immersion objective lens (LUMPlanFL N; Olympus). The microscope was placed on the antivibration table (Micro-g; TMC, Peabody, MA). To record an oscillation and displacement of hair bundles, the specimen was illuminated with a 630 nm light-emitting diode (UHP-T-SR; Prizmatix, Holon, Israel). The image was projected at a magnification of 1300X onto a dual photodiode, whose calibrated signal represented hair-bundle displacement.

### Estimation of hair-bundle stiffness

To measure the stiffness of hair bundles, force was applied by a flexible glass fiber (Figure S1A). After a quartz capillary 1.2 mm in diameter (Q120-60-7.5; Sutter Instruments, Novato, CA, USA) had been tapered with an electrode puller (P-2000; Sutter Instrument), its tip was melted and pulled laterally to form a shaft for hair-bundle stimulation. To enhance contrast on the photodiode, the fiber was sputter-coated with gold-palladium (Hummer 6.2; Anatech, Sparks, NV, USA). The stiffness and drag coefficient were estimated by measuring Brownian motion in water with photodiode sampling at 10,000 s^-1^ (25,26). The stiffness of the fibers (K_f_) was 150-400 μN·m^-1^.

A fiber mounted on a piezoelectric actuator (PA 4/12; Piezosystem Jena, Hopedale, MA, USA) was attached to a hair bundle at the kinociliary bulb with a micromanipulator (MP-285, Sutter Instrument). The fiber’s displacement (ΔX_f_) was controlled with LabVIEW 2019 software (National Instruments, Austin, TX, USA). The glass probe was displaced in five steps, ranging from 20 nm to 100 nm, to move the hair bundle in the negative direction such that the transduction channels were largely closed (Figure S1A, left and middle panels). Stimuli 100 ms in duration were presented every 200 ms. The values of ten repetitions of each stimulus magnitude were averaged and recorded. The hair-bundle displacement (ΔX_HB_) was calculated as the average value 30-45 ms after the onset of a stimulus pulse (Figure S1A). Finally, the stiffness of hair bundles of each step (K_HB (single step)_) was obtained by the following formula; K_HB (single step)_ = ((ΔX_f_ - ΔX_HB_)/ΔX_HB_)·K_f_. K_HB (single step)_ of each 5 step was averaged and used as stiffness of hair bundles (K_HB_). To measure the stiffness of hair bundles with broken tip links, we added 2 mM EDTA in endolymph, then displaced bundles as mentioned above (Figure S2A,B). Furthermore, to observe the displacement of the cuticular plate, a borosilicate glass microspheres 2 μm in diameter (Duke Scientific, Palo Alto, CA, USA) were placed on the apical cellular surface above the cuticular plate (Figure S3A) and the displacement of the glass beads was measured while the hair bundle was displaced (Figure S3B,C).

### Observation of spontaneous hair-bundle oscillation

Hair-bundle oscillation is readily observable with the optical setting described above. To quantify the displacements, 10 s segments of spontaneous hair-bundle oscillation were detected by the photodiode and recorded (LabVIEW 2019). A high-speed camera (Phantom VEO710L; Vision Research, Wayne, NJ, USA) was also used to record spontaneous oscillation. The video data were extracted as consecutive TIF images and the movement of hair bundles was tracked by the TrackMate plugin (27) of Fiji (ImageJ) (28). Oscillations were recorded before and after treatment with inhibitors and after washout.

### Recording of microphonic potentials

For the recording of microphonic potentials, which provide a means of assessing the average response of an ensemble of hair cells, a frog’s saccule was placed in a two-compartment experimental chamber (29,30). The saccule was sealed with cyanoacrylic adhesive (PeriAcryl 90) across a hole 500-700 μm in diameter through a circular plastic coverslip 10 mm in diameter and 200 μm in thickness (12547, Thermo Fisher Scientific). The mounted saccule was placed between gaskets of thermoplastic material (Parafilm, Amcor, Zurich, Switzerland) to provide electrical isolation and prevent the mixing of solutions. The coverslip was placed over the lower chamber, which was filled with artificial perilymph containing 99 mM Na^+^, 19 mM K^+^, 2 mM Ca^2+^, 118 mM Cl^-^, 3 mM **D**-glucose, and 5 mM HEPES at pH 7.3. By partially depolarizing the hair cells, this high-K^+^ perilymph minimized voltage-, time-, and Ca^2+^-sensitive currents and thus simplified the analysis of microphonic responses (29). The upper compartment was then added to the chamber and filled with artificial endolymph.

To elicit responses from a group of hair cells with similarly oriented hair bundles, we partially removed the otolithic membrane (29). Displacements produced by a piezoelectric actuator (PA 4/12, Piezosystem Jena) were applied to the remaining otolithic membrane by means of a glass probe produced by melting and roughening the tip of a micropipette. The displacement of the glass probe was controlled by LabVIEW 2019 software and the maximal amplitude was adjusted to the value that evoked a saturating response. The glass probe was displaced by ten positive and ten negative steps with equal separations in their magnitudes; an unstimulated control was also recorded. The responses were averaged across three repetitions. To estimate the resting open probability, we first determined the maximal positive response, *V*_100%_, and the greatest negative response, *V*_0%_, that were recorded between 10 ms and 20 ms after the onset of stimulation (Figure 4A). The resting open probability *P*_OR_ was then calculated as *P*_OR_ = -*V*_0%_ / (*V*_100%_ - *V*_0%_).

### Measurement of auditory brainstem responses

To assess the effect of inhibitors on hearing, we performed transtympanic injection of blockers prior to recording of ABRs from mice. Each animal was anesthetized with isoflurane delivered through a face mask at a concentration of 3 % - 5 % during induction and of 1 % - 2 % to maintain a surgical plane of anesthesia. The tympanic membrane was visualized under a microscope and incised with a 30-gauge needle (Figure 5A). After 5 μL of inhibitor solution or a saline control solution had been gently injected into the middle ear (31), the animal was allowed to awake and kept on a heating pad for 2 hr.

After a mouse had been anesthetized with an intraperitoneal injection of 100 mg/kg ketamine and 10 mg/kg xylazine, the ABR was recorded (32). Computer-generated stimuli were amplified by a stereo amplifier (SA1, Tucker-Davis Technologies, Alachua, FL, USA) and delivered by a free-field speaker (MF1, Tucker-Davis Technologies). Tone-burst stimuli with rise and fall times of 1 ms and durations of 10 ms were delivered 1000 times apiece at frequencies of 4 kHz, 8 kHz, 16 kHz, and 32 kHz. Subcutaneous electrodes were inserted in three positions: a recording electrode behind the pinna, a reference electrode in the scalp, and a ground electrode in the back near the tail. The electrical signals were amplified 10,000X and bandpass filtered between 300 Hz and 3000 Hz by a preamplifier (P55; Grass Instruments, West Warwick, RI, USA). The averaged responses were reviewed by two authors, R.O. and B.F., and hearing thresholds were determined by their agreement.

## Results

### Expression of NM2 and RLC in the frog’s saccule

Immunofluorescent labeling of the apical surfaces of the bullfrog’s saccular hair and supporting cells demonstrated NM2A, with MYH9 heavy chains, and NM2B, with MYH10 heavy chains (Figure 1A). These proteins occurred in a circumferential ring around the cuticular plate, but not in the hair bundle. Although absent from the apical surfaces, NM2C with MYH14 heavy chains occupied the apical cytoplasm of supporting cells. MYL12 was evident in hair bundles and MYL9 occurred at the apical surfaces of hair cells (Figure 1B). The apical surface of a frog saccular hair cell is therefore bounded by an actomyosin cable composed of NM2A and NM2B with MYL9 as the associated RLC (Figure 1C). Furthermore, MYL12 is expressed in the stereocilia independently of NM2.

**Figure 1.**
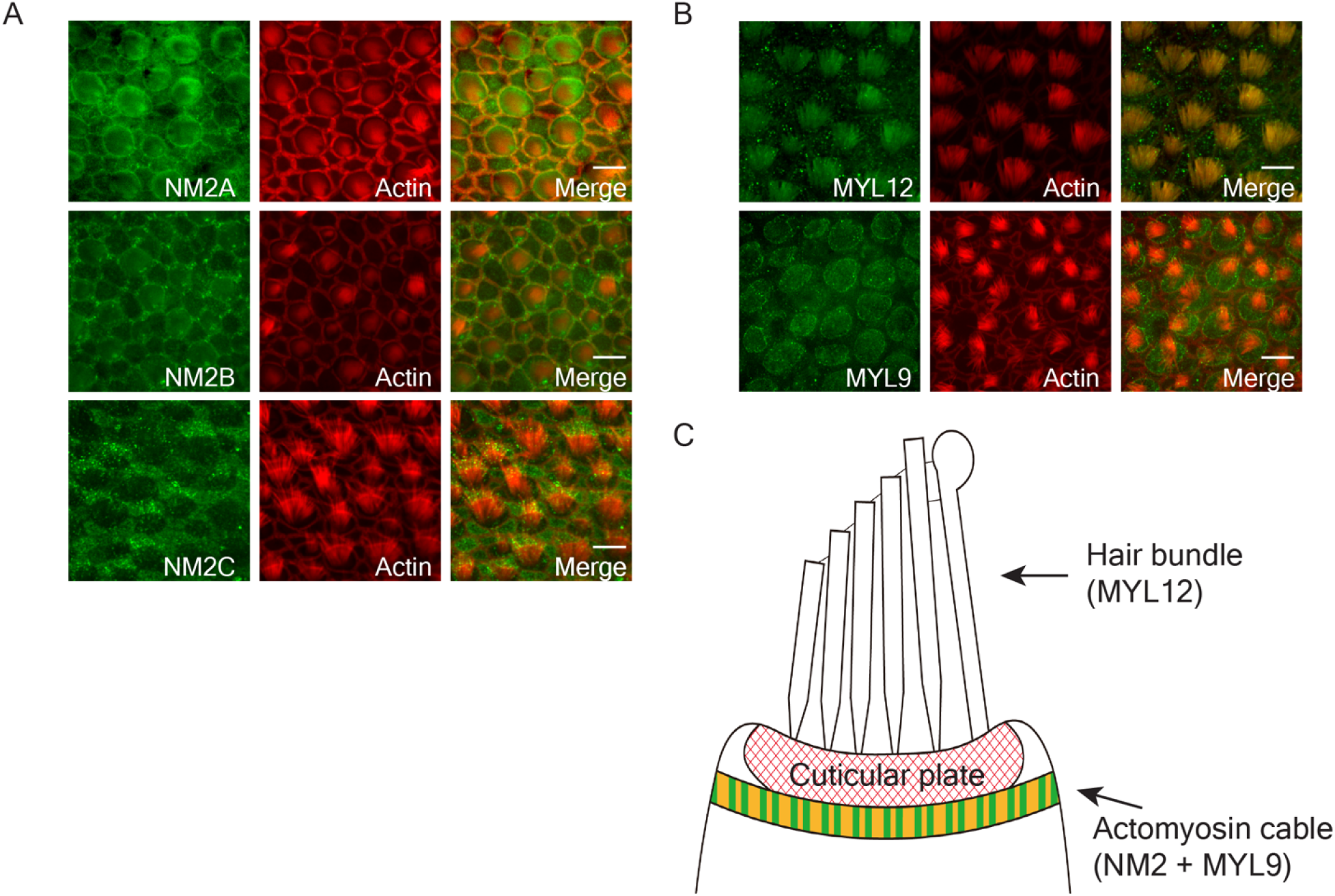
The expression of NM2 and myosin light chains in frog saccular hair cells. (A) Immunofluorescence labeling demonstrates NM2A and NM2B in the apical portions of hair cells and NM2C in supporting cells. Here and in panel *B*, actin labeled with phalloidin marks hair bundles, cuticular plates, and cellular perimeters. Scale bars here and in panel *B*, 10 μm. (B) Immunofluorescence labeling marks MYL12 in hair bundles and MYL9 in the apical cytoplasm of hair cells. (C) In a schematic diagram of the apical portion of a frog’s saccular hair cell, MYL12 is expressed in the hair bundles and non-muscle myosins and MYL9 are expressed in the actomyosin cable.

### Effect of inhibiting NM2 and MLCK on hair-bundle stiffness

To examine the physiological function of NM2 and MLCK, we assessed the effect of their inhibitors on the stiffness of hair bundles of the frog’s saccule (Figure S1A). Based on the results of control experiments, we performed stiffness measurement at room temperature (Figure S1B,C).

We treated the saccule with 10 μM blebbistatin, a nonspecific inhibitor of NM2 isoforms. Within 60 min, the stiffness declined significantly to 30 % of its control value (*P* = 0.0013, *N* = 10; Figure 2A,B). The effect was reversible: by 60 min after washout of the compound, the stiffness was not significantly different from the control value (*P* = 0.13, *N* = 10). Because blebbistatin significantly reduced the stiffness of hair bundles with disrupted tip links, the effect did not require intact transduction (Figure S2A).

**Figure 2.**
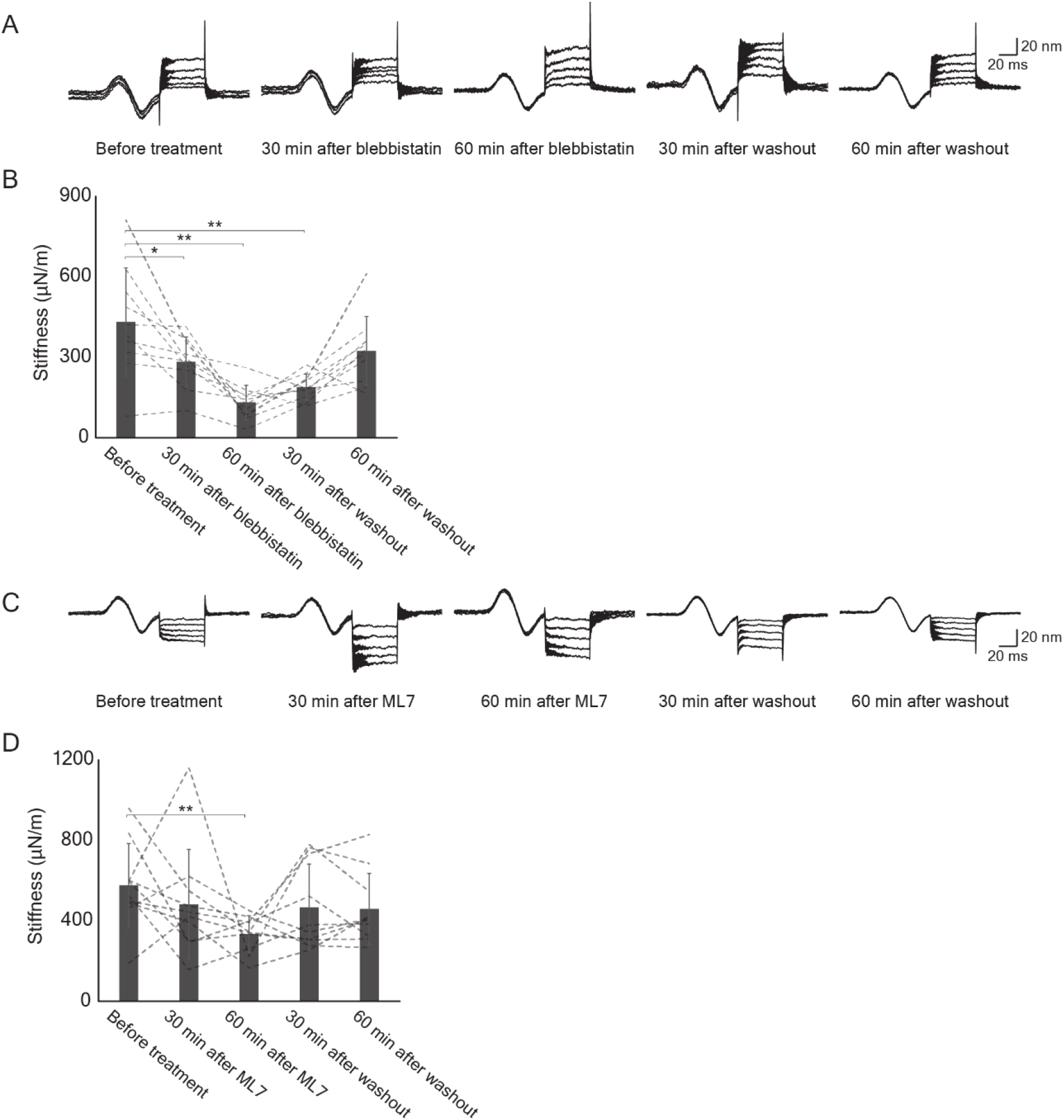
The effects of inhibitors on the stiffness of frog saccular hair bundles. (A) Records from a hair bundle before, during, and after treatment with 10 μM blebbistatin portray the average responses from ten repetitions of mechanical pulses of five positive amplitudes. Here and in panel *C*, the sinusoidal curves at the outset of each record represent displacements of the photodiode and serve as calibrations of ±20 nm (B) Averaged over ten repetitions, the data from panel *A* indicate that blebbistatin significantly and reversibly reduced bundle stiffness. Here and in subsequent figures: *, *P* < 0.05; **, *P* < 0.01. Note that there was no significant difference between the control records and those acquired after 60 min of washout. (C) Records from a hair bundle before, during, and after treatment with 2 μM ML7 typify the average responses from ten repetitions of mechanical pulses of five negative amplitudes. (D) Statistical analysis of the data from panel *C* demonstrates a significant difference between control and treated conditions but no difference 30 min or 60 min after washout.

At a concentration of 2 μM, ML7, an inhibitor of MLCK, also significantly affected the stiffness of saccular hair bundles. Treatment for 60 min lowered the stiffness to 58 % of its control level (*P* = 0.0068, *N* = 10; Figure 2C,D). The effect was reversible within 30 min after washout (*P* = 0.40, *N* = 10). In addition, ML7 reduced the stiffness of hair bundles with disrupted tip links; the effect, however, was not statistically significant (Figure S2B).

We measured the movement of the hair-cell surface during stimulation to confirm that the stability of the cuticular plate was not affected by inhibitors (Figure S3A). Glass beads placed on the cellular apex above the cuticular plate did not show significant displacements associated with stimulation (Figure S3B,C). Taken in conjunction with the foregoing result, the data imply that NM2 and RLC phosphorylation at the hair cell’s apex contribute to the hair bundle’s stiffness.

### Effect of inhibiting MLCK on spontaneous hair-bundle oscillation

When freed of their connections with an otolithic membrane and maintained in a two-compartment experimental environment, hair bundles of the frog’s saccule exhibit robust, low-frequency oscillations that reflect the active process of mechanoelectrical transduction (33) (Movies S1 and S4). To examine whether the spontaneous oscillation of hair bundles is affected by NM2 and MLCK, we used high-speed video microscopy or photodiode recording to observe spontaneous oscillations during inhibition of the two proteins. Because temperature affects the frequency and magnitude of oscillation, we performed control experiments to establish 20 °C as a suitable level for long-term observations (Figure S4A).

As has been observed previously (34), hair cells showed oscillations with a variety of frequencies and waveforms (Figure 3). Treatment of active hair bundles with 10 μM blebbistatin did not arrest oscillations, an observation that accords with evidence that the slow adaptation process of hair bundles involves MYO1C and MYO7A rather than NM2. The treatment did alter the characteristics of some oscillations; for example, an oscillation that initially occurred at a frequency near 3 Hz became biphasic and declined to a frequency of 1.5 Hz (Figure 3A).

**Figure 3.**
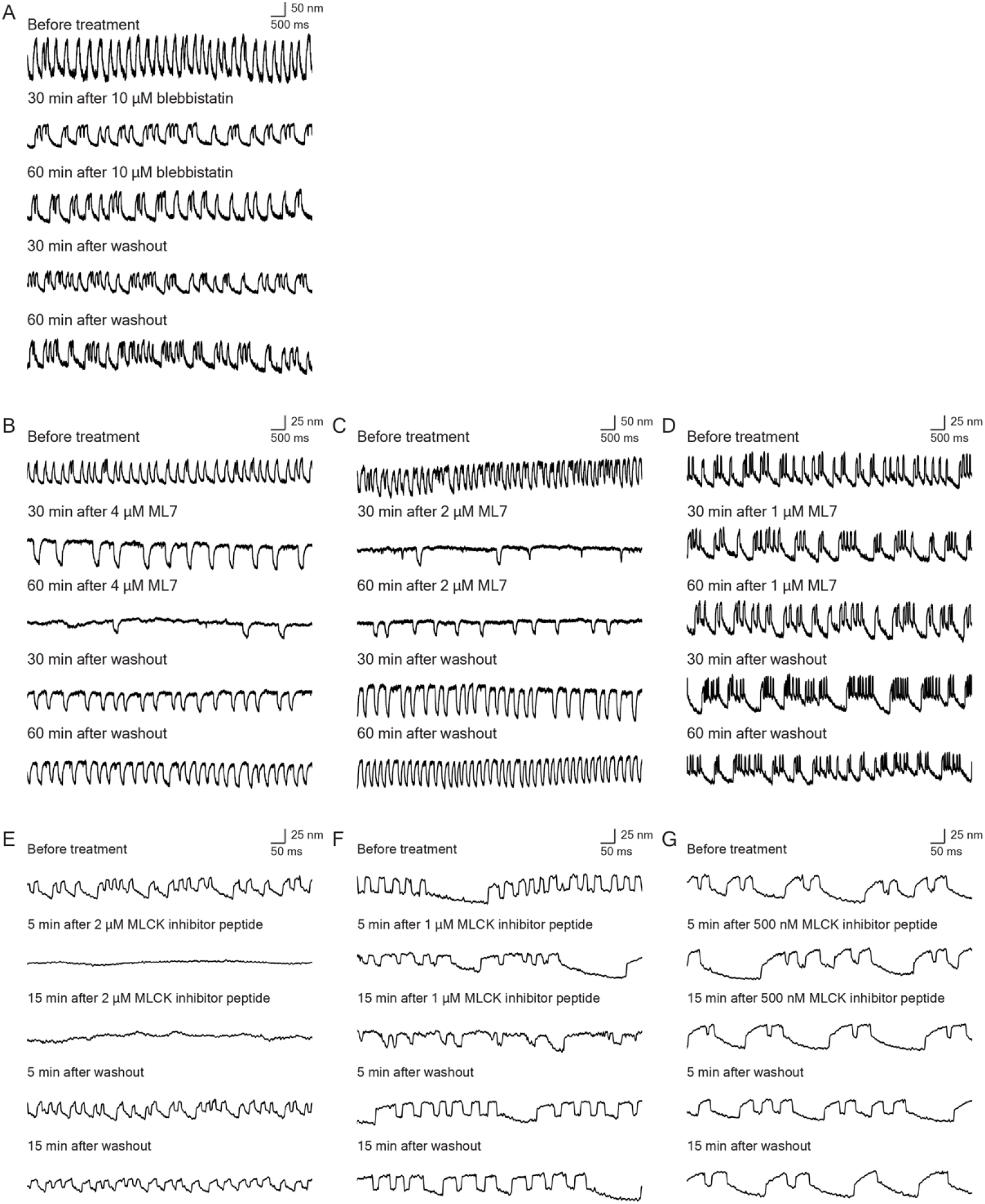
The effects of inhibitors on spontaneous oscillation by frog saccular hair bundles. (A) Treatment with 10 μM blebbistatin reversibly reduced the frequency of oscillation. (B) Treatment with 4 μM ML7 greatly slowed and almost arrested spontaneous oscillation; the effect was partially reversible after washout. (C) ML7 at a concentration of 2 μM had a similar but weaker effect than the foregoing; washout led to essentially complete recovery. (D) The effect of 1 μM ML7 was marginal. (E) Only 5 min of treatment with 2 μM MCK inhibitor peptide 18 completely blocked spontaneous oscillation, which largely recovered upon washout. (F) At a concentration of 1 μM, peptide 18 elicited a slight slowing of oscillations. (G) The effect of 0.5 μM peptide 18 was negligible. In all panels, an upward deflection signifies movement in the positive direction, towards the hair bundle’s tall edge.

We also observed the effect of ML7 on spontaneous oscillation. The motion was completely inhibited at a concentration of 2 μM or 4 μM, but showed recovery after washout (Figure 3B,C). At 1 μM, the compound did not significantly affect oscillation (Figure 3D). High-speed video microscopy confirmed the photodiode observations (Movies S1-S3; Figure S4B). To test an alternative means of inhibition, we also used MLCK inhibitor peptide 18. At a concentration of 1 μM or 2 μM, oscillation was again inhibited and recovered after washout (Figure 3E,F); an inhibitor concentration of 500 nM had no effect (Figure 3G). High-speed video imaging confirmed the inhibition of oscillation by 2 μM of the peptide (Movies S4-S6; Figure S4C). Inhibition by peptide 18 was more rapid than that owing to ML7. Both of the MLCK inhibitors arrested hair bundles during displacement in the positive direction, that is, while deflected toward their tall edges.

### Effect of inhibiting MLCK on transduction channels’ open probability

We next inquired whether inhibiting NM2 or MLCK would affect mechanoelectrical transduction in frog saccular hair cells. To investigate the roles of the proteins in adjusting the resting open probability of mechanoelectrical-transduction channels, we applied inhibitors while recording microphonic responses (29). These extracellular signals reflect the total current that traverses transduction channels and provide a convenient means of assessing the averaged responses of hundreds of hair cells (Figure 4A).

**Figure 4.**
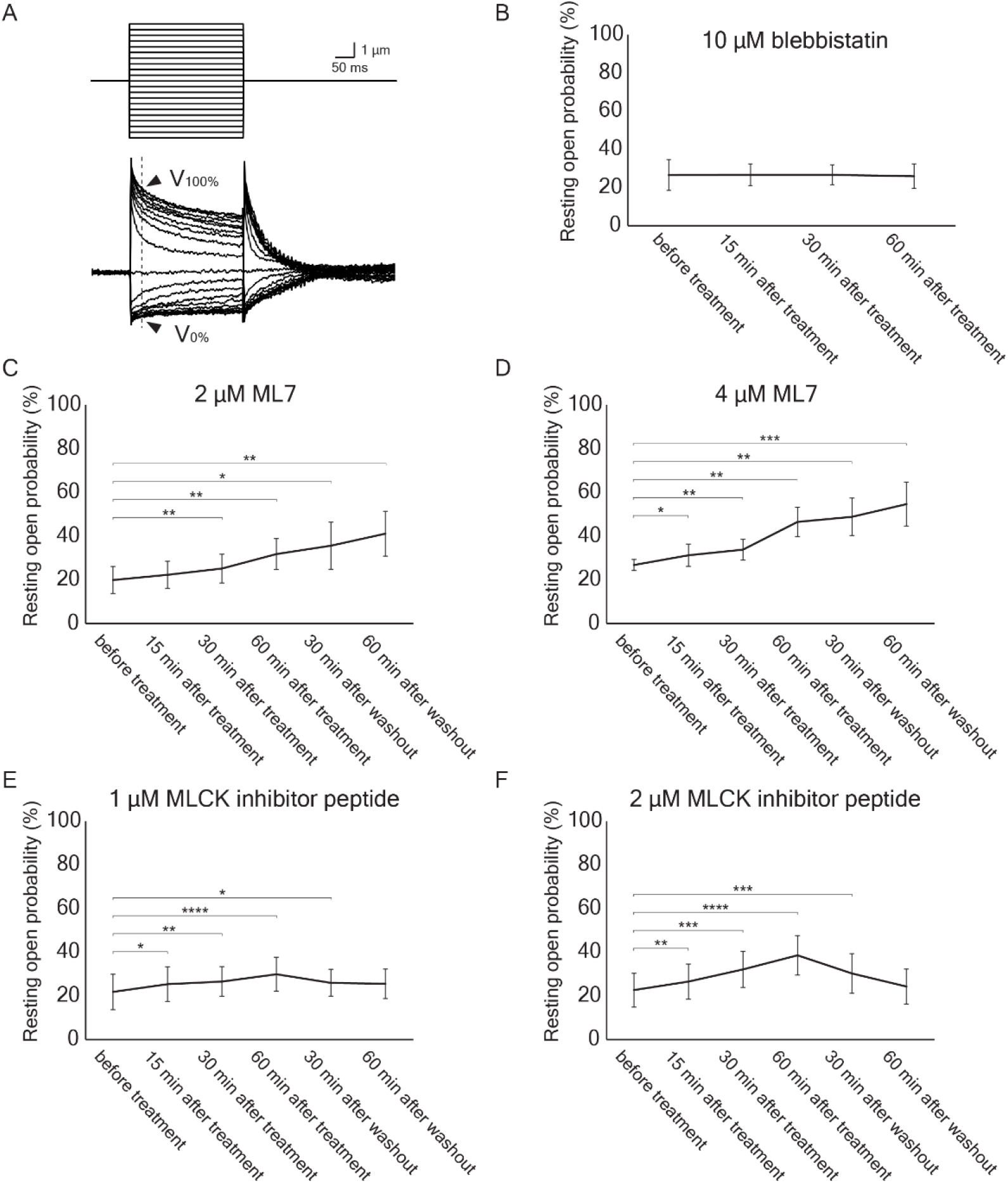
The effects of inhibitors on the resting open probability of transduction channels. (A) In recordings of microphonic responses, a glass probe displaced the otolithic membrane to 21 levels, including zero (upper traces). The maximal response (*V*_100%_) and minimal response (*V*_0%_) were estimated from averaged electrical records (arrowheads) 10 ms after after the onset of stimulation (lower traces). (B) Treatment with 10 μM blebbistatin had no effect on the resting open probability. (C) In six experiments, exposure to 2 μM ML7 slightly increased the resting open probability, but the effect was not reversed after washout. (D) Treatment with 4 μM ML7 in six experiments had a greater but irreversible effect. (E) At a concentration of 1 μM, MLCK inhibitor peptide 18 significantly increased the resting open probability. (F) In six experiments, 2 μM MLCK inhibitor peptide 18 significantly and reversibly raised the resting open probability. There was no significant difference between the control recording and that 60 min after washout. Here and in the subsequent figure: ***, *P* < 0.001; ****, *P* < 0.0001.

Consistent with the expectation that tip-link tension is set by MYO1C and MYO7A, treatment with the NM2 inhibitor blebbistatin did not significantly affect the resting open probability (Figure 4B). By contrast, 30 min of treatment with 2 μM of the MLCK inhibitor ML7 raised the resting open probability significantly, from 19.7 ± 6.2 % to 25.0 ± 6.6 % (*P* = 0.009, *N* = 6; Figure 4C). The effect was not reversed by 60 min of washout of the inhibitor. The resting open probability also showed a significant and lasting elevation after treatment with 4 μM ML7 (Figure 4D).

Peptide 18 also had a significant effect on the resting open probability of transduction channels. After 60 min of treatment with 1 μM of peptide 18, the probability increased from 21.6 ± 6.8 % to 29.7 ± 7.8 % (*P* < 0.0001, *N* = 6; Figure 4E). Following 60 min of washout, the open probability of 25.3 ± 6.8 % did not differ significantly from the control value (*P* = 0.086). Furthermore, treatment with 2 μM of peptide 18 evoked a significant increase of the resting open probability and a washout effect as well (Figure 4F). This and the foregoing experiment indicate that MLCK inhibition causes a significant increase in open probability.

### Effect of MLCK inhibition on the murine auditory brainstem response

To investigate the effect of MLCK inhibition on hearing, we measured the threshold of the auditory brainstem response (ABR) after the transtympanic injection of ML7 of mice (Figure 5A). In keeping with the volume of the murine middle ear (31), 5 μL of ML7-containing solution was injected into the middle ear on one side and 5 μL of saline solution was injected into the middle ear on the opposite, control side.

**Figure 5.**
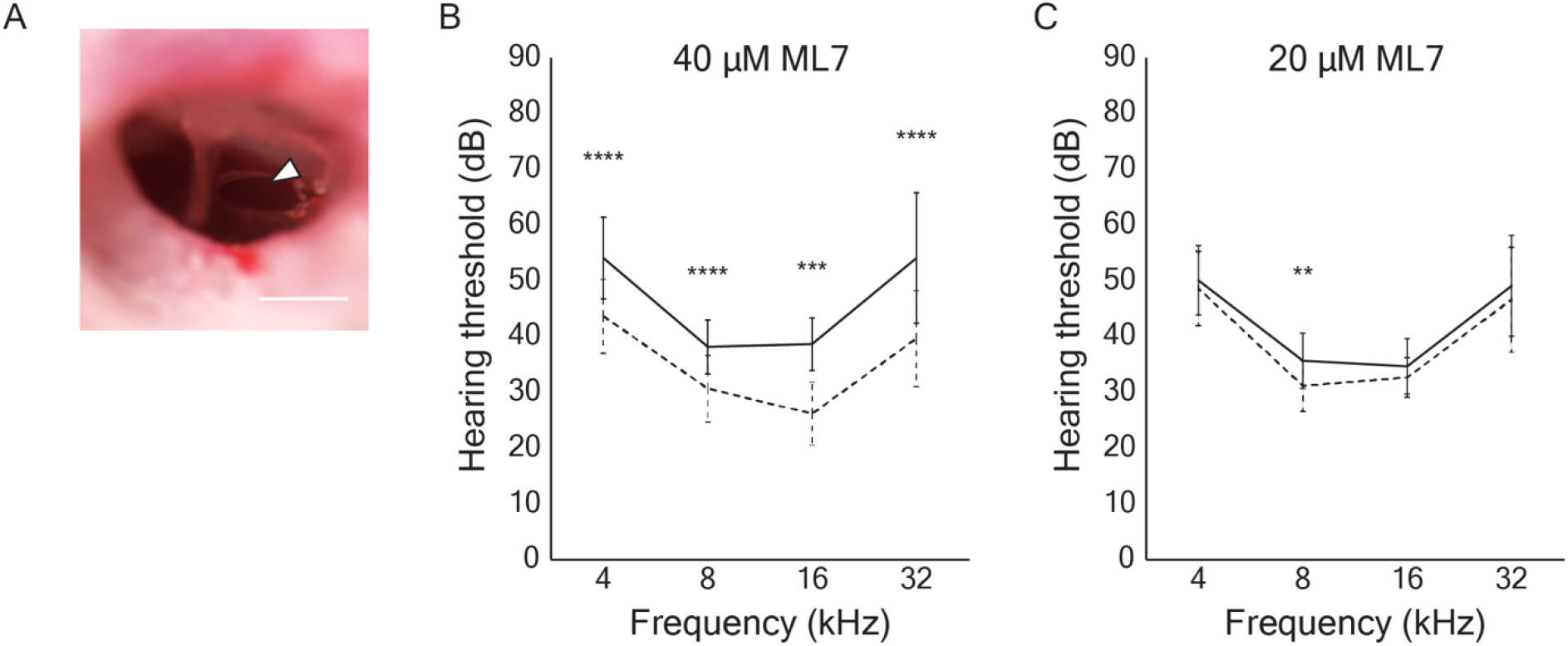
The effect of ML7 on hearing thresholds in the mouse. (A) In an image of the tympanic membrane after transtympanic injection, the white arrow indicates the incision of the tympanic membrane for drug administration. Scale bar, 0.5 mm. (B) Following the administration of 40 μM ML7 into the middle ear, averaged auditory brainstem responses displayed significantly elevated thresholds at the four frequencies tested. The solid line marks responses from the ear exposed to the inhibitor, whereas the dotted line shows data for the control ear exposed to vehicle. (C) A lower concentration of ML7, 20 μM, caused a significant threshold elevation only at a stimulus frequency of 8 kHz.

Two hours after transtympanic injection at a concentration of 40 μM, ML7 significantly increased the ABR threshold at stimulus frequencies from 4 kHz to 32 kHz (Figure 5B). A lower concentration of the compound, 20 μM, yielded a significant threshold elevation only at 8 kHz (Figure 5C).

### Effect of inhibition of MLCK on phosphorylation of RLC

We performed immunolabeling to ascertain whether the phosphorylation of RLC was significantly affected by MLCK inhibitors. Using frog saccules from microphonic recordings after 60 min of treatment with 4 μM ML7, we found that immunolabeling of phosphorylated RLC disappeared from hair bundles but not from the apical surfaces of hair cells (Figure 6A). Immunolabeling of phosphorylated RLC was also performed in the cochleas of mice after ABR recordings. In animals treated with 40 μM ML7 the immunofluorescence signal for phosphorylated RLC was far weaker in hair bundles on the treated than the untreated side of an animal (Figure 6B). These control experiments demonstrate that ML7 treatment inhibited the phosphorylation of RLC in both species and support the inference that the observed physiological effects stemmed from reduced RLC activity.

**Figure 6.**
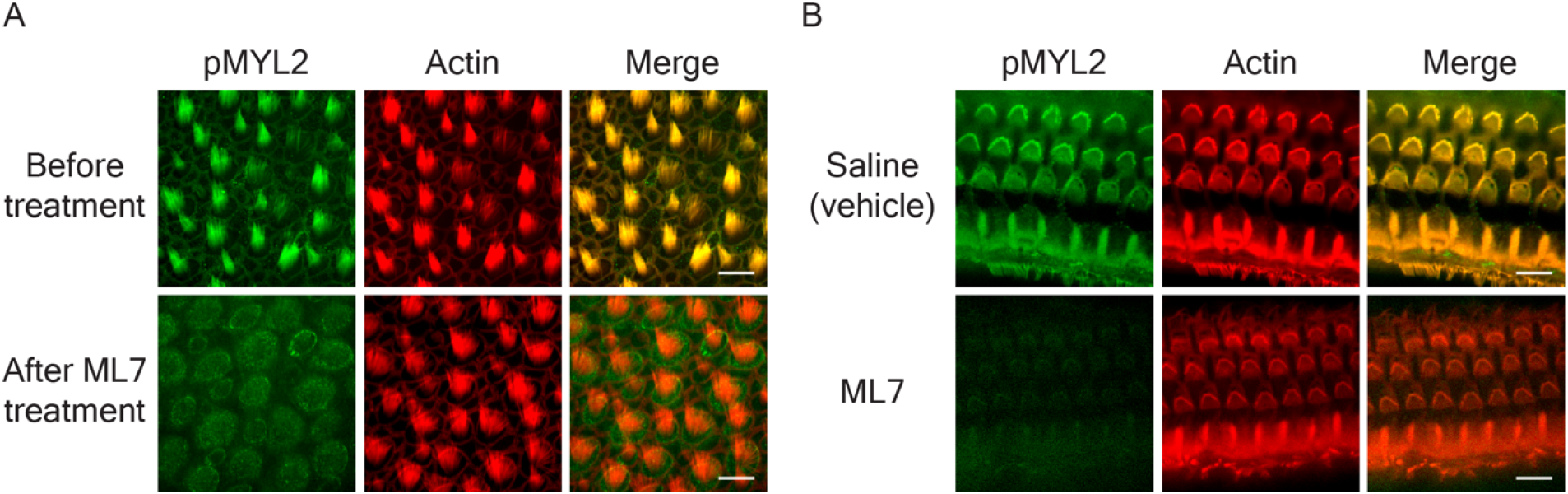
The effect of ML7 on the phosphorylation of RLC in frog and mouse hair bundles. (A) Immunofluorescence labeling of the phosphorylated regulatory light chain pMYL2 shows a robust signal in frog hair bundles that is abolished by treatment with 4 μM ML7. Scale bar, 10 μm. (B) In the mouse organ of Corti, immunofluorescence labeling of phosphorylated light chain is abolished after the transtympanic injection 40 μM ML7. In this instance the control specimen reflects the result after transtympanic injection of the saline vehicle. Scale bar, 10 μm.

## Discussion

This study of the physiological function of NM2 and MLCK in inner ear hair cells has yielded several results. Frog saccular HCs express NM2 and MYL9 in the apical surface and MYL12 in the hair bundles. The bundles’ stiffness declines after treatment with blebbistatin or ML7. Moreover, spontaneous oscillation of the frog saccular HCs is slightly affected by treatment with blebbistatin and is inhibited by MLCK inhibitors such as ML7 and MLCK inhibitor peptide 18. The resting open probability of frog saccular HCs was increased by treatment with ML7 or peptide 18. Finally, transtympanic injection of ML7 in mice middle ear increases the threshold of auditory brainstem responses.

Although the bullfrog has been used extensively in physiological research, it has played less of a role in biochemical investigations. NM2 and RLC have been shown to occur at the apical surfaces of hair cells in rodents (11,12). In the current study, we determined that frog saccular hair cells express NM2 and MYL9 at their apical surfaces and MYL12 in their hair bundles. Our measurements indicate that NM2 plays a role in regulating the stiffness of frog saccular hair bundles. Because we displaced bundles in the negative direction that reduced the tension in tip links, the results bear primarily on the stiffness of the stereociliary pivots. Our other control studies also showed that inhibitors decreased the hair-bundle stiffness even after tip links had been broken by EDTA treatment (Figure S2A,B). This result showed that the inhibition of NM2 and MLCK affected the stiffness of stereociliary pivots, which reflects the anchorage of stereociliary rootlets into the cuticular plate. In conclusion, NM2 and MLCK activity affect the tightness of the insertion of stereociliary rootlets into the cuticular plate or indeed the stiffness of the cuticular plate itself.

The spontaneous oscillation of hair bundles results from the active process of non-mammalian vertebrates (24). Extensive evidence suggests that oscillation emerges from the activity of myosin motors at the tops of hair bundles (35). In the present study, we found that spontaneous oscillation is inhibited by the MLCK inhibitors ML7 and peptide 18. Because peptide 18 is membrane permeable, it acts quickly and its inhibitory effect is reversible. Although the two inhibitors have distinct inhibitory mechanisms (36,37), their similar and concentration-dependent effects are consistent with MLCK inhibition. The results suggest that RLC serves as a light chain of MYO1C, perhaps by binding to that isoform’s highly conserved second IQ domain (38).

We observed that the resting open probability of transduction channels increases when MLCK is inhibited, presumably because phosphorylation of MYL12 reduces tip-link tension and facilitates channels’ closure. MYL12 binds to unconventional myosins including MYO15A (39) and MYO7A, a myosin motor of cochlear hair cells (40). Both of these unconventional myosins possess IQ motifs to which their light chains bind (22). MYO1C also has IQ motifs that bind calmodulin (41) and could potentially bind light chains such as MYL12. We therefore hypothesize that MLCK phosphorylates MYL12 bound to MYO1C in the frog’s hair bundles, and that inhibition of this phosphorylation reduces the sliding of MYO1C along the actin filaments of stereociliary cores. Consistent with this hypothesis, MLCK inhibitors arrest spontaneously oscillating hair bundles with a deflection in the positive direction. We hypothesize that a complex of MYO1C and MYL12 is necessary to produce force during adaptation and that the phosphorylation of MYL12 enhances this force production (Figure 7).

**Figure 7.**
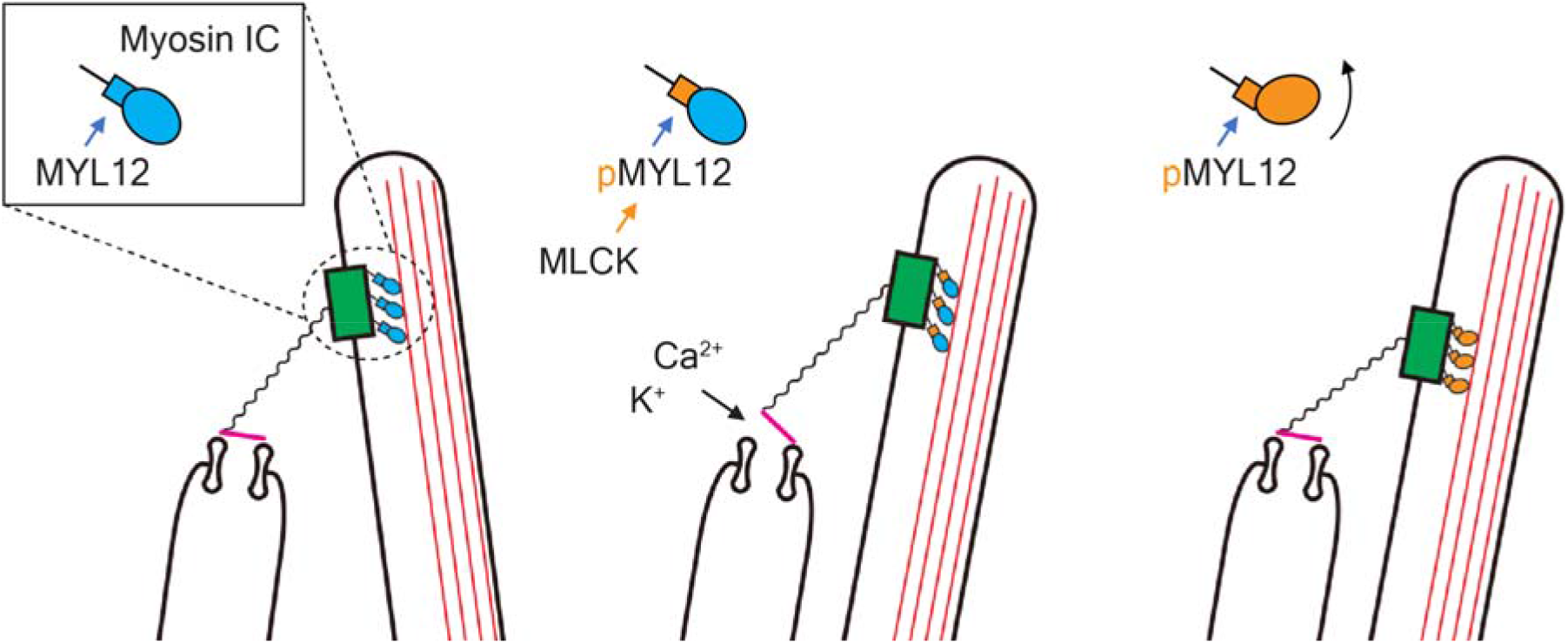
The hypothesized effect of inhibitors on the adaptation motor of hair bundles. The left schematic diagram shows the control configuration of a hair bundle, with the bundle in its resting position and the transduction channels largely closed. In the middle diagram, deflection of the bundle in the positive direction opens transduction channels, which admit K^+^ and Ca^2+^ into the cell. As shown in the right diagram, phosphorylation of the regulatory light chain MYL12 promotes slippage of the myosin motor along stereociliary actin filaments, which presumably slackens tip links and promotes the consequent reclosure of channels.

In addition to our *ex vivo* experiments, we showed that hearing in intact mammals is inhibited by ML7. The experiments demonstrated dose dependency and effects over a broad range of stimulus frequencies. Previous evidence also supports a role of MLCK activity in hearing: mice exhibiting a specific deletion of MLCK in either inner or outer hair cells display increased ABR thresholds (19,20). The phosphorylation of MYL12 is therefore important in normal hearing, perhaps by affecting an adaptation motor such as MYO7A. However, there are two limitations in this experiment. First, we do not know the exact concentration of ML7 in the inner ear. ML7 presumably gains access to the cochlea through the round window, but the efficacy of this infiltration is unknown. The inhibitory constant of ML7 for MLCK is 300 nM (42), so concentrations of 20-40 μM might inhibit other kinases as well. The second reservation is that we did not observe a consistent effect of peptide 18 on the ABR. Because the peptide is highly cell-permeable and easy to wash out, it is difficult to maintain an effective concentration during protracted recordings.

## Statistics

Statistical analyses were performed using JMP version 17 statistical software (IBM Japan, Tokyo, Japan). All data are expressed as means ± standard deviations. A paired Student’s *t*-test was performed for the comparison; a value of *P* < 0.05 was considered to represent a significant difference.

## Data availability

The data generated and analyzed during the current study are available from the corresponding author upon reasonable request.

## Author Contributions

R.O. and A.J.H designed the study and wrote the manuscript. R.O. and B.F. performed the experiments. K.W. wrote Python programs for the analysis of microphonic measurements. R.G.A. participated in frog dissections and in measurements of hair-bundle stiffness and spontaneous oscillation.

## Acknowledgements

The authors thank Leslie Diaz for advice on mouse experiments and members of our group for comments on the manuscript. K.W. received a fellowship from Weill Cornell Medical College. R.O., B.F., and R.G.A. were supported by Howard Hughes Medical Institute, of which A.J.H. is an Investigator.

