## Supplemental information for "Influence of myosin regulatory light chain and myosin light chain kinase on hair cells of the inner ear"

Contents:

Supplementary Results

Supplementary Figures and Legends

**Supplementary Results**

**Optimization of experimental temperature**

Because the temperature possibly affects the stiffness of hair bundles, we first measured the stiffness art various temperatures (Figure S1A‑C). At higher temperatures including 30 ℃ and 35 ℃, the stiffness exceeded that at 20 ℃ or 25 ℃ (Figure S1B,C). The stiffness at 35 ℃ (1235 ± 975 μN·m^‑1^) was significantly greater than those at the other three temperatures (378 ± 215 μN·m^‑1^ at 20 ℃, *P* = 0.011; 327 ± 149 μN·m^‑1^ at 25 ℃, *P* = 0.011; 356 ± 174 μN·m^‑1^ at 30 ℃, *P* = 0.021; Fig. S1C). Because temperature was not critical for the stiffness at temperatures below 35 ℃, further measurements were performed at room temperature.

**Stiffness measurement with broken tip links**

To confirm that blebbistatin and ML7 affected the stiffness without the tension of tip links, we measured hair-bundle stiffness with tip links broken by the addition of 2 mM EDTA to the endolymph. After breakage of tip links, the stiffness dropped to 25 % of the control value (Figure S2A,B). Furthermore, in six experiments subsequent 10 μM blebbstatin treatment further reduced the stiffness from 276 ± 164 μN·m^‑1^ after treatment with 2 mM EDTA to only 88.3 ± 48.7 μN·m^‑1^ (Figure S2A; *P* = 0.0068). In six experiments, 2 μM ML7 treatment also caused a decrease in stiffness from 250 ± 239 μN·m^‑1^ after treatment with 2 mM EDTA to only 85 ± 54 μN·m^‑1^; however, the difference was not statistically significant (Figure S2B; *P* = 0.20).

**Control for cuticular-plate movement**

When the hair bundle is pushed or pulled, the stereocilia should pivot upon the cuticular plate of a hair cell. However, there is an alternative possibility that the cuticular plate is displaced or depressed. To examine this issue, we placed the glass beads on the apical cellular surface above the cuticular plate and measured the movement of glass beads by pushing or pulling the hair bundle (Figure S3A). Because the beads were stuck to the cellular surface, movement of the glass beads reflected that of the cuticular plate. After treatment with 10 μM blebbistatin or 2 μM ML7, the glass beads did not show significant movement corresponding to the displacement of hair bundles (Figure S3B,C). We concluded that blebbistatin and ML7 exerted their effects on hair bundles or the structure to which the stereocilia are anchored rather than on the stability of the subjacent cuticular plates.

**Supplementary Figures and Legends**

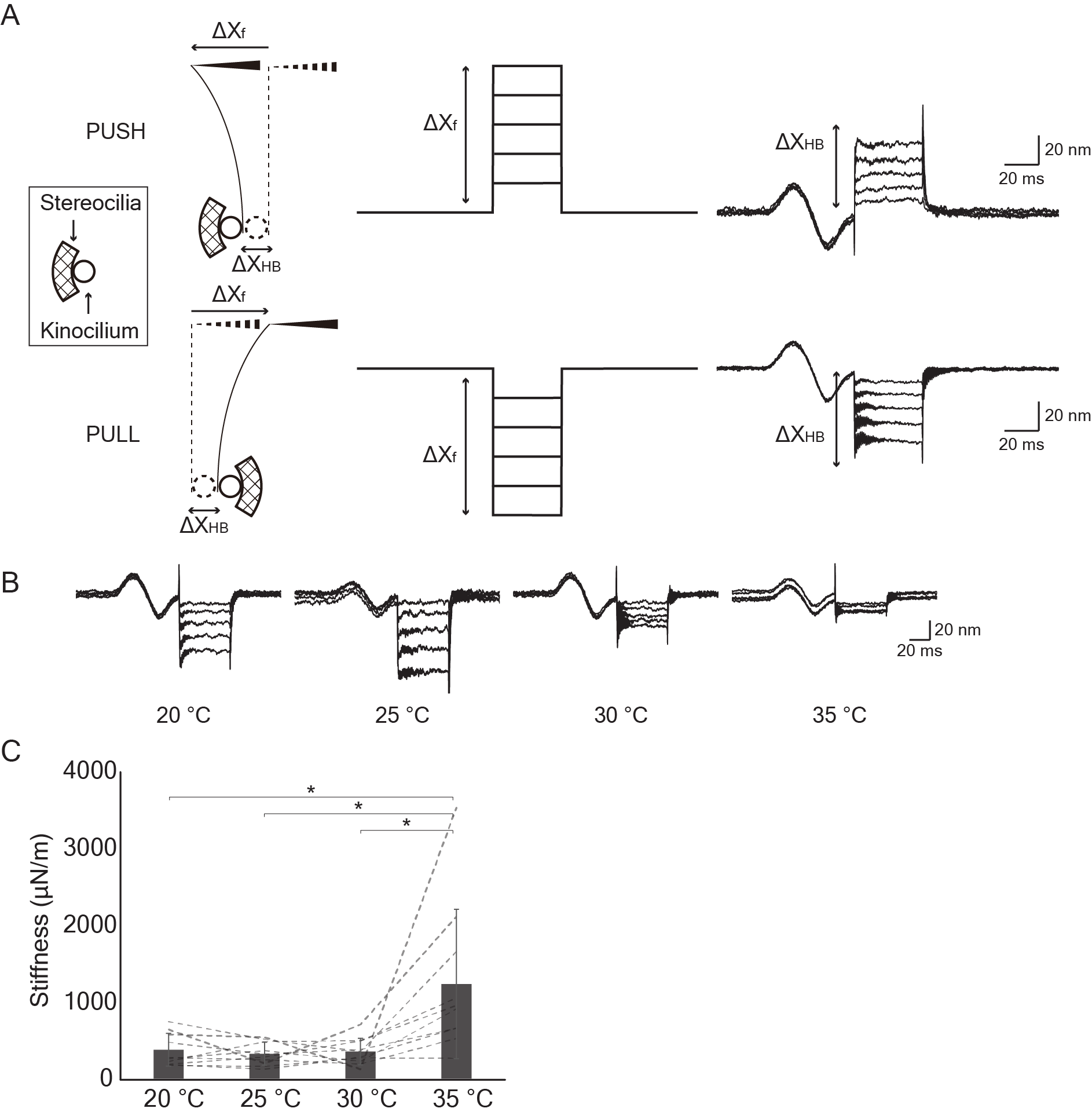

**Figure S1. Measurement of the stiffness of a frog saccular hair bundle.** (A) The schematic drawing (left panel) shows the positioning of oppositely oriented hair bundles as they are deflected by a flexible stimulus fiber. The movement of the fiber's base (middle panels) is induced by a piezoelectrical stimulator controlled by a computer program. The actual responses of a hair bundle (right panels) follow a biphasic calibration signal of ±20 nm. (B)The response of hair bundle at various temperatures shows progressive stiffening between 25 ˚C and 35 ˚C. (C) The data from panel *B* show that the stiffness at 35 ℃ significantly exceeds that at lower temperatures.

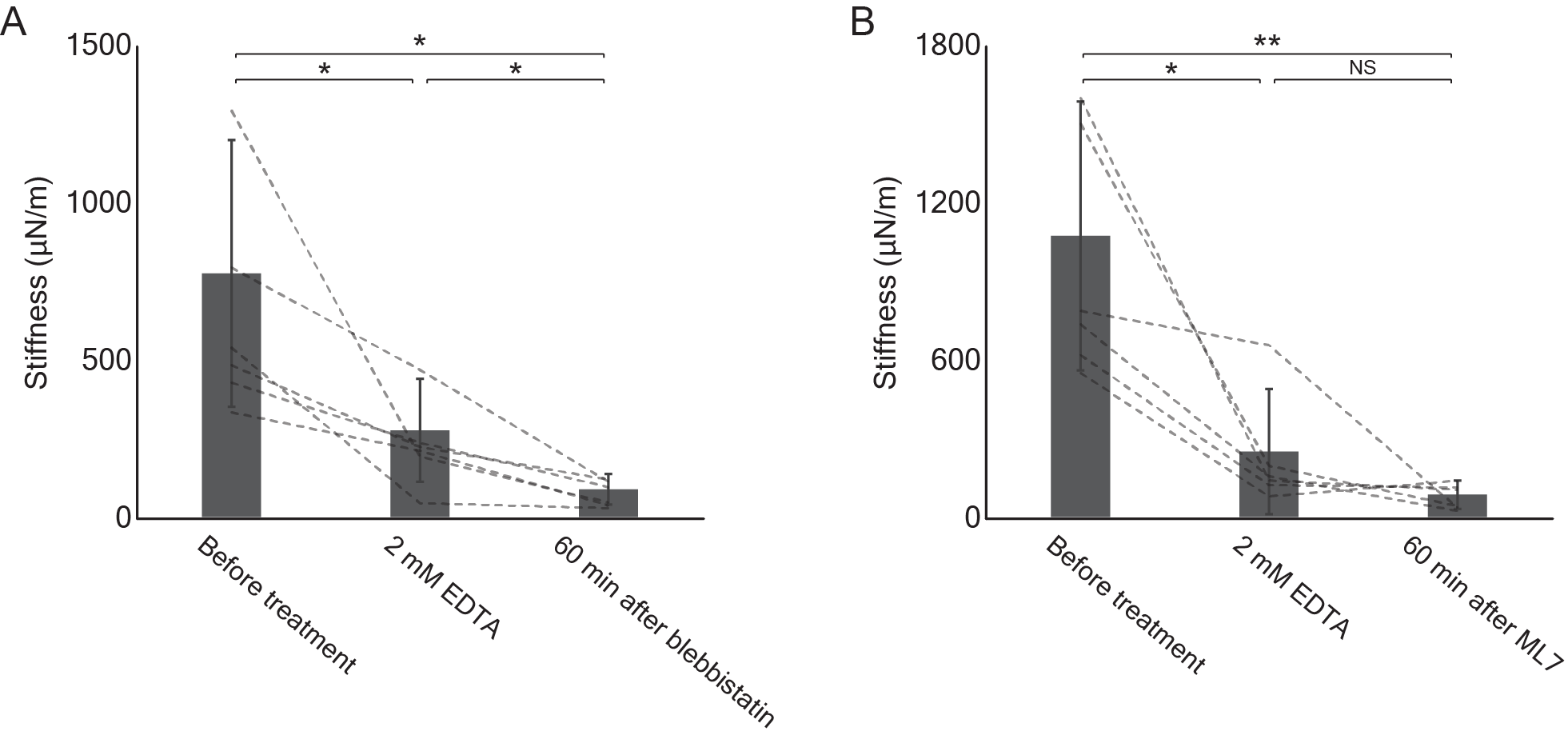

**Figure S2. The effects of inhibitors on the stiffness of frog saccular hair bundles with broken tip links.** (A) After the tip links had been broken by exposure to 2 mM EDTA in endolymph, hair-bundle stiffness declined significantly in six experiments. Treatment with blebbistatin evoked a significant additional reduction. (B) In six experiments, treatment with ML7 reduced the stiffness of hair bundles with broken tip links. In this instance, however, the difference was not significant.

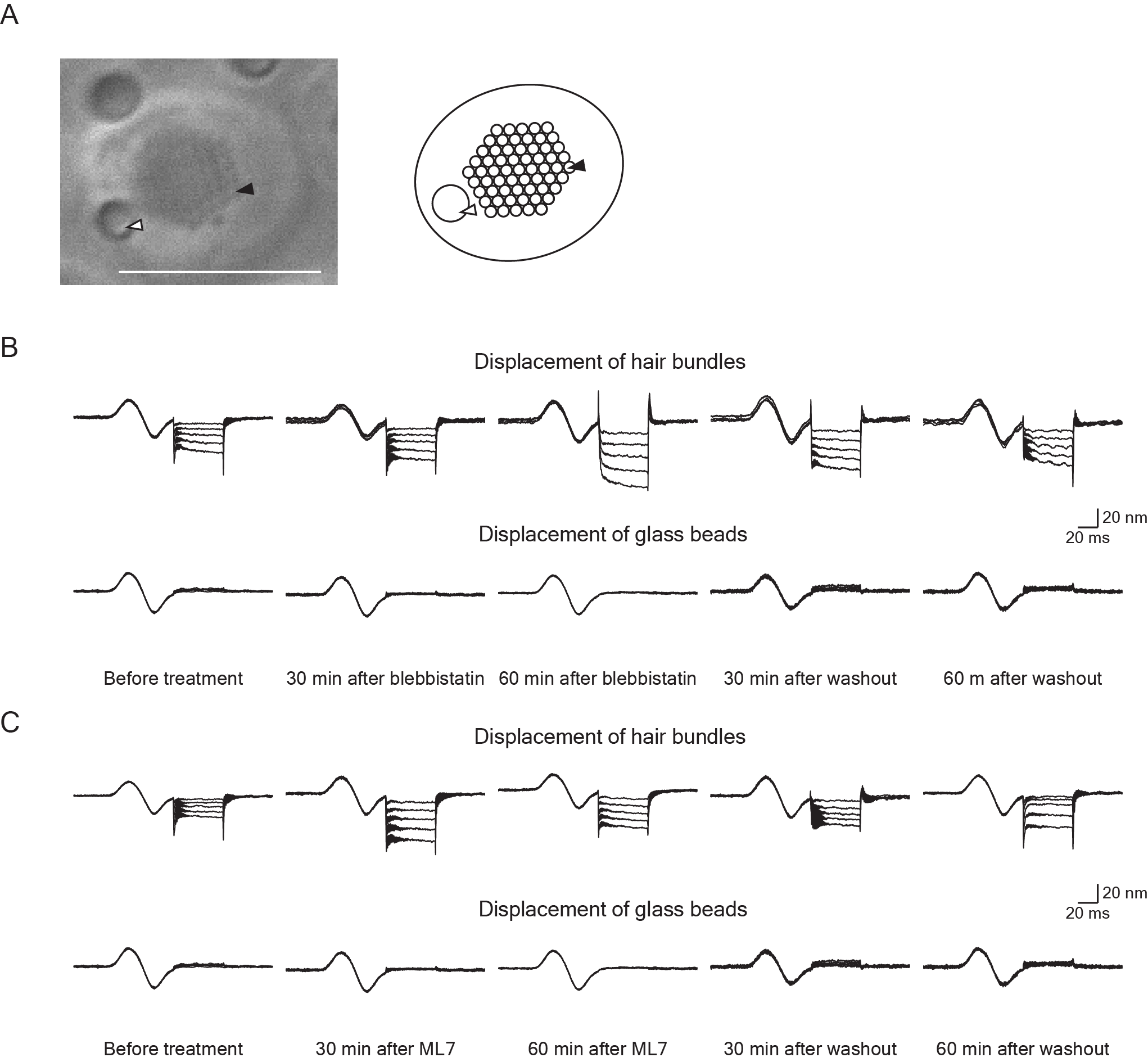

**Figure S3. Measurement of apical-surface displacements during stiffness measurements.** (A) A microscopic image (left panel) and schematic drawing (right panel) show glass beads in the apical surface of a hair cell immediately above the cuticular plate. In each image, a glass bead is indicated by the white arrowhead and the cluster of stereocila by the black arrowhead. Scale bar, 10 μm. (B) Glass beads did not show significant displacements during stiffness measurement with 10 μM blebbistatin (C) During exposure of a hair bundle 2 μM ML7, glass beads also showed no significant displacement.

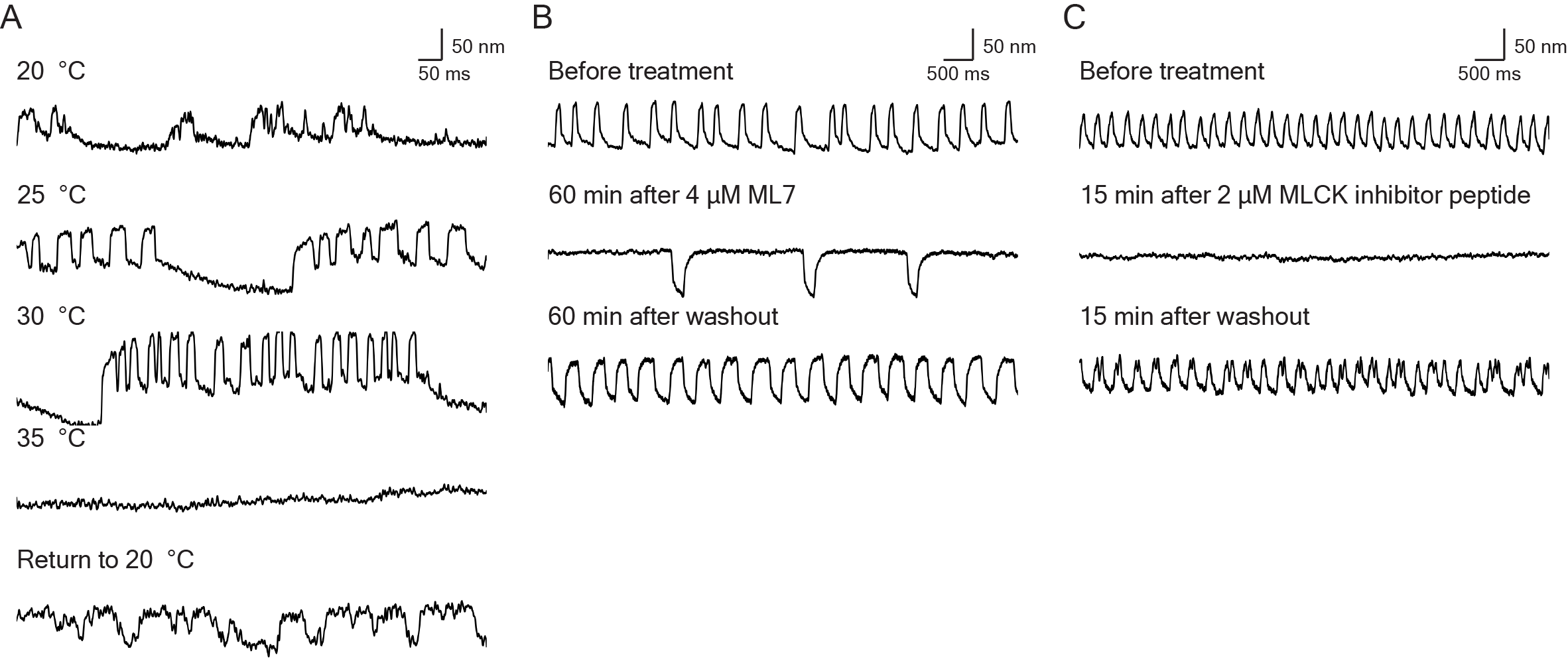

**Figure S4. Observation of spontaneous oscillation by frog saccular hair bundles.** (A) Oscillation became faster at higher temperatures until it was arrested at 35 ˚C. (B) A high-speed camera recorded the waveforms of spontaneous oscillation before and during treatment with 4 μM ML7 as well as after washout. These data correspond to Movies S1, S2, and S3. (C) The waveforms of the spontaneous oscillation recorded by a high-speed camera show the inhibitory effect of treatment with 2 μM MLCK inhibitor peptide 18. These data correspond to Movies S4, S5, and S6. In all panels, an upward deflection signifies movement in the positive direction, towards the hair bundle's tall edge.

**Supplementary Movies and Legends**

**Movie S1. Spontaneous oscillation of bullfrog saccular hair bundles before treatment recorded with a high-speed camera.** For this and the other Supplementary Movies, the scan rate was 1000 frames per second and the duration was 8 s.

**Movie S2. Spontaneous oscillation of bullfrog saccular hair bundles 60 min after treatment with 4 μM ML7.**

**Movie S3. Spontaneous oscillation of bullfrog saccular hair bundles 60 min after washout of ML7.**

**Movie S4. Spontaneous oscillation of bullfrog saccular hair bundles before treatment with peptide 18.**

**Movie S5. Spontaneous oscillation of bullfrog saccular hair bundles 15 min after treatment with 2 μM inhibitor peptide 18.**

**Movie S6. Spontaneous oscillation of bullfrog saccular hair bundles 15 min after washout of peptide 18.**
